# Calcium Dynamics During Pollen Tube Reception in *Arabidopsis* Ovules

**DOI:** 10.64898/2026.03.30.715275

**Authors:** Chiharu Kato, Teruaki Goromaru, Takuya T. Nagae, Yoko Mizuta, Daisuke Kurihara, Yoshikatsu Sato, Satohiro Okuda, Tetsuya Higashiyama

**Affiliations:** Department of Biological Sciences, Graduate School of Science, The University of Tokyo, Hongo, Bunkyo-ku, Tokyo 113-0033, Japan; Division of Biological Science, Graduate School of Science, Nagoya University, Furo-cho, Chikusa-ku, Nagoya, Aichi 464-8602, Japan; RIKEN Center for Sustainable Resource Science (CSRS), Suehiro-cho, Tsurumi, Yokohama, Kanagawa 230-0045, Japan; Institute for Advanced Research (IAR), Nagoya University, Furo-cho, Chikusa-ku, Nagoya, Aichi 464-8601, Japan; Institute of Transformative Bio-Molecules (WPI-ITbM), Nagoya University, Furo-cho, Chikusa-ku, Nagoya, Aichi 464-8601, Japan

**Keywords:** *Arabidopsis thaliana*, Ca^2+^ dynamics, N-linked glycosylation, Pollen tube reception, Sporophytic tissues in the ovule, Synergid cells

## Abstract

In flowering plants, pollen tubes communicate with ovular cells to achieve precise one-to-one pollen tube reception. The final step of this communication between the pollen tube and synergid cells has been extensively investigated and visualized by calcium imaging. Synergid cells exhibit characteristic cytoplasmic calcium concentration oscillations, which are thought to play a critical role in pollen tube reception. However, their significance and relationship with calcium dynamics in the entire ovule remain unclear. Here, we show, using the calcium sensor GCaMP6s, that proteins involved in asparagine-linked glycosylation (N-linked glycosylation) are required for normal calcium oscillations in synergid cells but are not essential for pollen tube reception. Using a semi-*in vivo* assay in *Arabidopsis thaliana*, we found that the amplitude of these oscillations prior to rapid pollen tube growth across the filiform apparatus was reduced in mutants lacking the oligosaccharyltransferase (OST) 3/6 subunit or alpha1,2-glucosyltransferase (ALG) 10, both of which are involved in N-linked glycosylation. Notably, these mutants did not exhibit reduced fertility attributable to defects in the female gametophyte but instead showed a polytubey phenotype due to a sporophytic defect. These findings suggest that N-linked glycans mediate communication between synergid cells and the pollen tube and indicate that the typical pattern of calcium oscillations in synergid cells is not essential for triggering pollen tube rupture. Furthermore, we show that sporophytic tissues of the ovule exhibit calcium waves that propagate toward the funiculus in correlation with pollen tube contact and rupture, implying that ovular tissues can potentially transmit these signals distantly beyond the ovule. Together, these findings reveal previously unrecognized intercellular calcium signaling and its significance in pollen tube reception by the ovule.

## Introduction

Interaction between male and female reproductive tissues is essential for successful sexual reproduction in both animals and plants, and calcium signaling regulates multiple steps in this process (Dresselhaus et al. 2016). In flowering plants (angiosperms), immotile sperm cells are delivered to the ovule by the tip-growing pollen tube (Dresselhaus et al. 2016). The pollen tube elongates within the pistil while undergoing multiple guidance and selection events and ultimately enters the embryo sac.

Upon contact between the pollen tube and the two synergid cells located at the micropylar entrance of the embryo sac, both the pollen tube and one synergid cell rupture, releasing the sperm cells into the embryo sac. This series of intercellular communications is called pollen tube reception, in which both male- and female-derived factors participate. In synergid cells, these include the receptor kinase FERONIA (Huck et al. 2003, Rotman et al. 2003, Escobar-Restrepo et al. 2007) and its GPI-anchored co-receptor LORELEI (Capron et al. 2008, Tsukamoto et al. 2010, Li et al. 2015, Liu et al. 2016), the receptor kinases ANJEA and HERCULES RECEPTOR KINASE 1 (HERK1) (Galindo-Trigo et al. 2020), the calcium channel NORTIA (Kessler et al. 2010) and the N-linked glycosylation–related proteins EVAN and TURAN (Lindner et al. 2015). In the pollen tube, key components include the transcription factors MYB97/101/120 (Leydon et al. 2013, Liang et al. 2013, Zhong et al. 2022), multiple RAPID ALKALINIZATION FACTOR (RALF) peptides that interact with FERONIA, ANJEA, HERK1 (Zhong et al. 2022) and the calcium pump ACA9 (Schiøtt et al. 2004). Loss of these factors leads to an overgrowth phenotype, in which the pollen tube fails to rupture and instead continues to elongate within the embryo sac, eventually coiling. ACA9 deficiency also causes an entry-stop phenotype, in which pollen tube growth ceases after contact with synergid cells (Schiøtt et al. 2004, Desnoyer et al. 2024). Pollen tube-expressed receptor kinases ANXUR1/2 (Boisson-Dernier et al. 2009) and BUPS (Zhu et al. 2018, Ge et al. 2017) have also been reported to be involved in pollen tube rupture through interactions with ovule-derived RALF peptides (Ge et al. 2017). Although the complete pathway remains unclear, pollen tube and synergid cell rupture is controlled by a multistep process involving coordinated signaling between male and female tissues.

During pollen tube reception, characteristic calcium oscillations have been observed in synergid cells (Iwano et al. 2012, Ngo et al. 2014, Denninger et al. 2014, Hamamura et al. 2014). Proteins expressed in synergid cells, including FERONIA, LORELEI and NORTIA (Ngo et al. 2014), as well as those expressed in the pollen tube, such as MYB97/101/120 (Ponvert and Johnson 2024), ACA9 (Desnoyer et al. 2024) and RALF4/19 (Gao et al. 2022), are involved in this process. These findings suggest that both female and male gametophytes contribute to the regulation of calcium dynamics in synergid cells. Moreover, loss-of-function mutants of these components exhibit a high frequency of failed pollen tube reception. Denninger et al. (2014) also reported that low-amplitude calcium oscillations do not trigger pollen tube rupture, and Duan et al. (2014) showed that calcium depletion in the pistil induces pollen tube overgrowth. Given the correlation between aberrant calcium oscillations and reduced fertility, calcium oscillations in synergid cells are thought to play a pivotal role in regulating pollen tube reception.

Ngo et al. (2014) classified synergid cell calcium dynamics into multiple phases based on pollen tube growth behavior. Upon contact with the pollen tube, cytosolic calcium oscillations are induced in synergid cells (Phase 1), during which pollen tube growth slows near the filiform apparatus (a thickened cell wall structure associated with the two synergid cells). In the subsequent phase (Phase 2), the pollen tube grew rapidly between the two synergid cells. Concurrently, cytosolic calcium levels in the receptive synergid cell remain elevated until pollen tube rupture and sperm discharge occur. These calcium dynamics reflect a highly coordinated communication between synergid cells and the pollen tube; however, their precise biological significance and underlying molecular mechanisms remain unclear.

Recent advances in live-cell imaging have revealed that sporophytic tissues of the ovule are also required for successful one-to-one pollen tube guidance and reception by the ovule (Mizuta et al. 2024). For example, when the first pollen tube starts to grow along the funiculus of the ovule, repulsion of a second pollen tube is initiated within 15 min and is sufficient to block polytubey by 45 min (Mizuta et al. 2024). These cell-to-cell communications occur before the tip of the first pollen tube arrives at the synergid cell, and sporophytic tissues contribute to these processes. However, calcium dynamics across the entire ovule, including sporophytic tissues, remain largely unexplored.

In this study, we monitored calcium dynamics in ovules under semi-in vivo conditions using GCaMP6s, a genetically encoded fluorescent calcium sensor characterized by high sensitivity and slow decay kinetics (Chen et al. 2013). This sensor has also been used in plant gametophytic cells (Gao et al. 2022, Matsuura-Tokita et al. 2026). We identified oligosaccharyltransferase 3/6 (OST3/6) and alpha-1,2-glucosyltransferase 10 (ALG10) as key regulators of synergid cell calcium oscillations. These proteins are involved in N-linked glycosylation (Farid et al. 2011, Farid et al. 2013). We also discovered that both the female gametophyte and sporophytic tissues of the ovule exhibited distinct calcium waves that coincided with pollen tube contact and rupture. Our findings reveal multilayered calcium signaling in both gametophytic and sporophytic tissues and provide a refined model for the regulation of pollen tube reception.

## Results

### Calcium oscillations in synergid cells are initiated upon tight contact between the pollen tube and the filiform apparatus

First, we observed calcium dynamics in synergid cells to validate the calcium sensor GCaMP6s during pollen tube reception. To monitor cytosolic calcium dynamics in synergid cells, the calcium sensor GCaMP6s was expressed under the control of the synergid cell-specific *MYB98* promoter (Kasahara et al. 2005). Pollen tube behavior was simultaneously monitored by expressing mApple (Shaner et al. 2008) under the control of the pollen-specific *LAT52* promoter (Twell et al. 1990). Consistent with a previous report (Ngo et al. 2014), we observed characteristic Phase 1 and Phase 2 calcium dynamics in synergid cells under semi-*in vivo* conditions (top graph in Fig. 1A), correlating with pollen tube growth behavior (bottom graph in Fig. 1A). There were variations in the pattern, duration, and initiation timing of Phase 1 calcium oscillations; however, the shift to Phase 2, as indicated by an increase in pollen tube growth speed, was consistently observed and was accompanied by a sustained higher level of calcium concentration prior to pollen tube rupture (Fig. 1 and Supplementary Fig. S1).

**Fig. 1.**
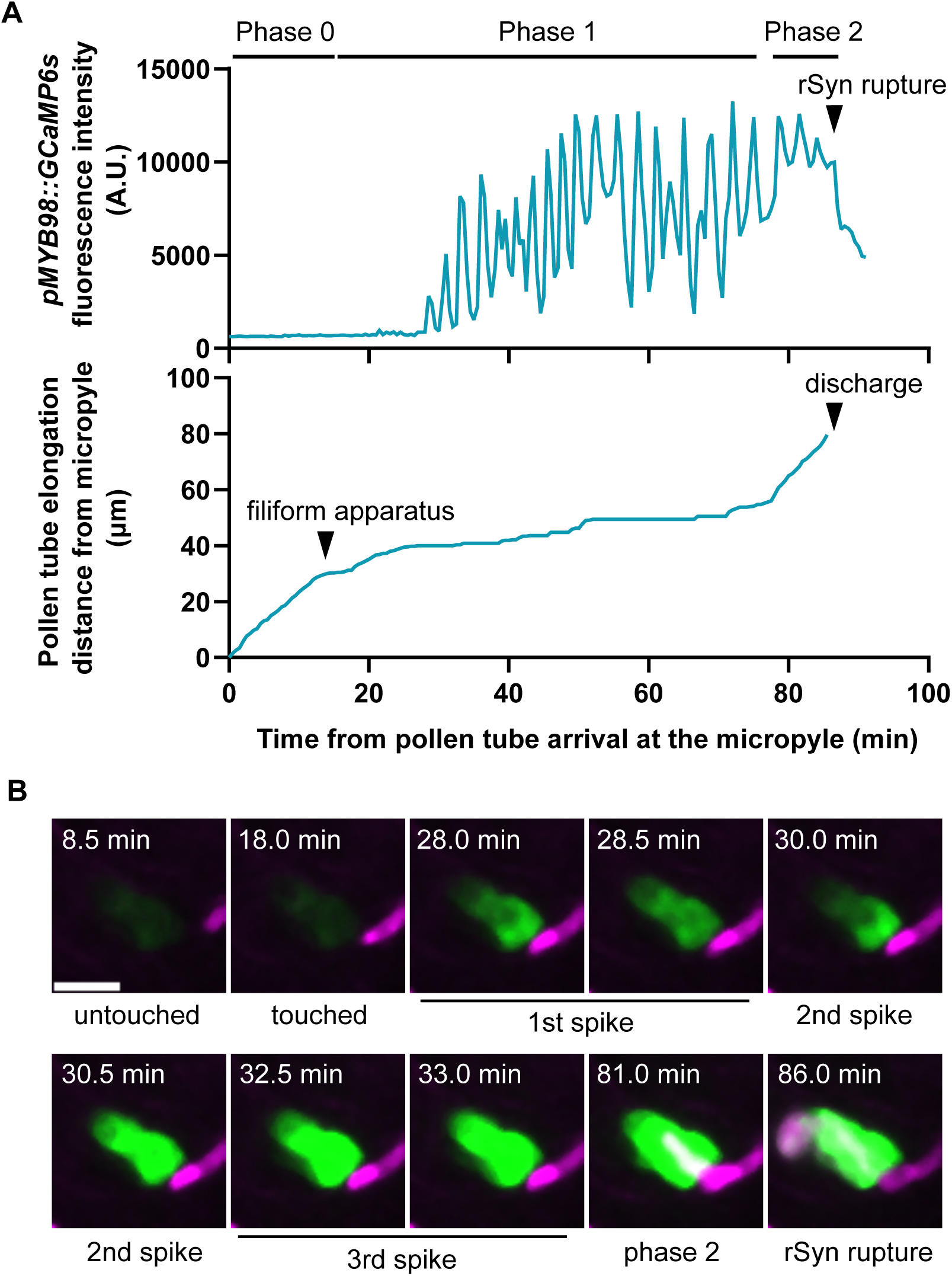
Wild-type synergid cells exhibit phase changes in cytoplasmic calcium ion concentration associated with pollen tube behavior during pollen tube reception. (A) Traces of the fluorescence intensity of GCaMP6s in synergid cells (top graph) and pollen tube elongation distance from the micropyle (bottom graph). The arrowhead labeled “filiform apparatus” shows when the pollen tube tip reaches the filiform apparatus, which marks the first point of physical contact between the pollen tube and synergid cells. rSyn, receptive synergid cell. (B) Time-lapse series of synergid cells expressing GCaMP6s (green) and a pollen tube expressing mApple (magenta), as measured in (A). The time (min) is the same as that shown in (A). Scale bar, 20 μm.

Compared with previously used calcium sensors, most of which were FRET-based ratiometric sensors (Denninger et al. 2014, Hamamura et al. 2014, Ngo et al. 2014, Ponvert and Johnson 2024), GCaMP6s appeared to be more sensitive, as expected (Chen et al. 2013), which was advantageous for monitoring the initiation of calcium oscillations. To capture the initiation of calcium oscillations in synergid cells, we defined the interval from when the pollen tube reached the micropyle to when it contacted the filiform apparatus as Phase 0. Under our conditions, most ovules did not show an increase in GCaMP6s fluorescence in synergid cells during Phase 0 (Fig. 1 and Supplementary Fig. S1).

In most ovules, calcium oscillations in synergid cells did not initiate even when the pollen tube reached the filiform apparatus (Fig. 1 and Supplementary Fig. S1). In these ovules, immediately after contact with the filiform apparatus, only a localized transient increase in calcium concentration was observed at the micropylar end of the synergid cells (Fig. 1 and Supplementary Fig. S1).

After the pollen tube made tight contact with the filiform apparatus, calcium waves initiated at the micropylar end of the synergid cells and propagated toward the chalazal pole (Fig. 1B). These waves were repeatedly generated, resulting in calcium oscillations (Fig. 1B). During Phase 1, the pollen tube slowly elongated along the filiform apparatus. In Phase 2, the GCaMP6s signal appeared nearly saturated and was observed as a sustained fluorescence level (Fig. 1 and Supplementary Fig. S1). The sharp spike upon pollen tube rupture, previously observed using YC3.60 (Ngo et al. 2014, Hamamura et al. 2014, Ponvert and Johnson 2024), was not clearly detected with GCaMP6s. Instead, a steep decrease in the signal was observed upon pollen tube discharge (Fig. 1 and Supplementary Fig. S1).

In summary, GCaMP6s was useful for time-lapse imaging of synergid cells during semi-*in vivo* pollen tube reception. The initiation of calcium oscillations was clearly observed with GCaMP6s and was triggered by tight contact between the pollen tube tip and the filiform apparatus.

### Loss-of-function mutants of the OST3/6 subunits and ALG10 involved in N-linked glycosylation show attenuated calcium oscillations in synergid cells

To dissect the molecular mechanism underlying calcium oscillations in synergid cells and the significance of their patterns, we searched for mutants defective in calcium oscillations. To minimize variation in GCaMP6s expression levels, each mutant was crossed with a wild-type line carrying *pMYB98::GCaMP6s*, which was used as a control. When we observed mutants related pollen tube reception using a semi-*in vivo* system, we found that loss-of-function mutants of OST3/6 subunit (*aru-1*) (Müller et al. 2016) and ALG10 (*alg10-1*) (Farid et al. 2011, Zupunski 2020) exhibited attenuated calcium oscillations in synergid cells during pollen tube reception, similar to *fenonia* (*fer-4*) mutant (Fig. 2; Supplementary Figs. S2–S5).

**Fig. 2.**
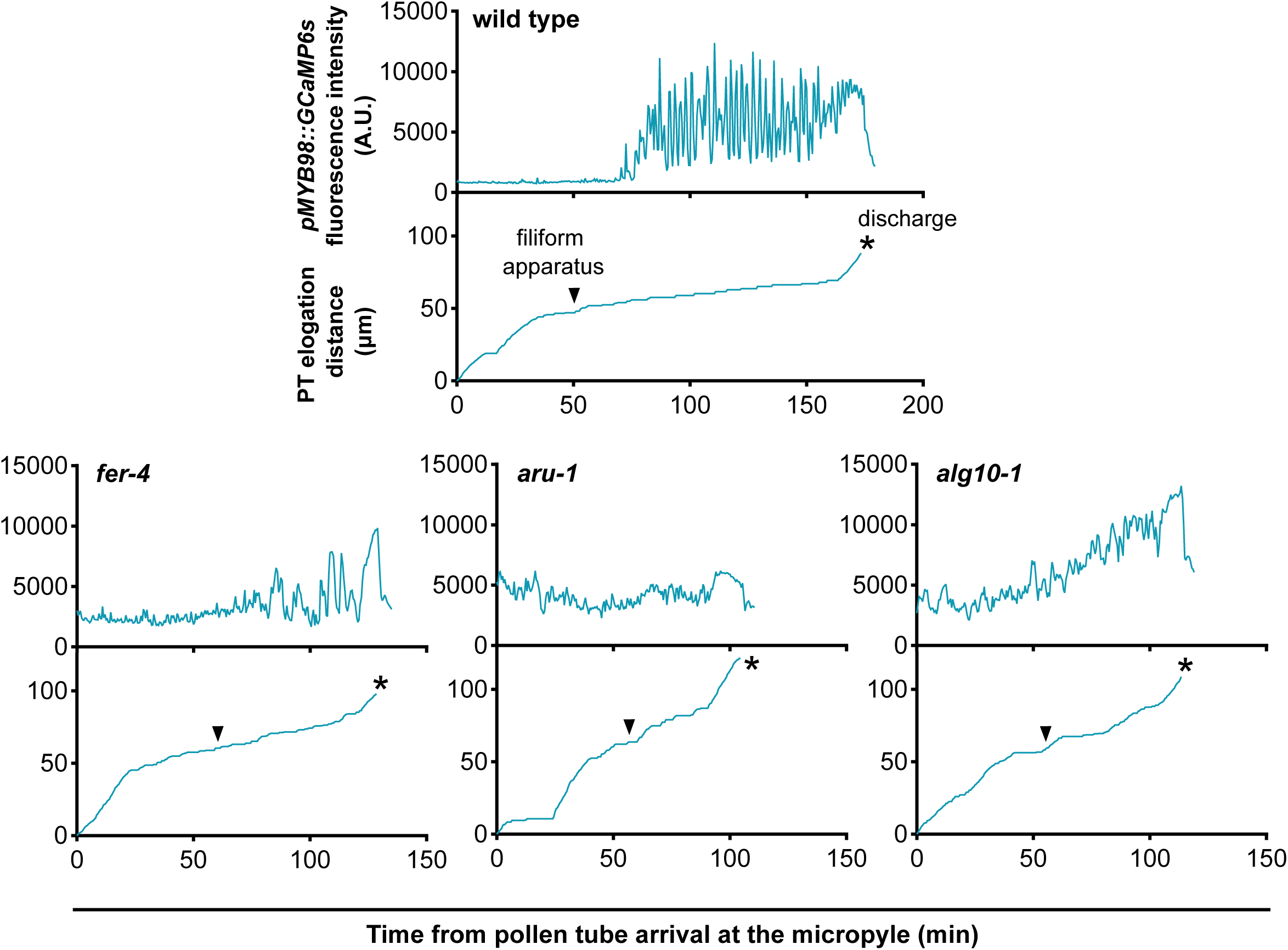
*aru-1* and *alg10-1* exhibit reduced amplitude of the synergid calcium oscillations during pollen tube reception. Each upper graph shows example traces of the fluorescence intensity of GCaMP6s in synergid cells, and the lower graphs show pollen tube elongation distance from the micropyle. Females with a wild type, *fer-4*, *aru-1,* or *alg10-1* background were pollinated with wild-type pollen. The arrowhead labeled “filiform apparatus” shows when the pollen tube tip reached the filiform apparatus. Asterisks indicate the time of pollen tube–synergid cell discharge. Pollen tube discharge was observed in some *fer-4* ovules under our semi-*in vivo* conditions.

OST3/6 and ALG10 are components of the N-linked glycosylation machinery (Farid et al. 2011, Farid et al. 2013), a major co-translational protein modification pathway (Supplementary Fig. S2). Many proteins localized to the cell surface or secreted into the extracellular space undergo N-linked glycosylation, and N-linked glycans play crucial roles in cell–cell interactions. Both genes act downstream pathway of EVAN and TURAN, which have been reported to be involved in pollen tube reception (Lindner et al. 2015). ALG10 localizes to the endoplasmic reticulum and functions as an α1,2-glucosyltransferase that adds a glucose residue to the terminus of lipid-linked oligosaccharides during N-glycan biosynthesis (Farid et al. 2011). These oligosaccharides are subsequently transferred to nascent polypeptides by the OST complex, which includes the OST3/6 subunit (reviewed in Aebi, 2013). Both genes are ubiquitously expressed, and mutants defective in either gene show a marked reduction in protein N-linked glycosylation.

In these mutants, arrival of the pollen tube at the filiform apparatus did not trigger a pronounced increase in cytosolic calcium levels in synergid cells (Fig. 2; Supplementary Figs. S3–S4). The amplitude of calcium oscillations was quantified by calculating the change in GCaMP6s intensity over a 30-second period, as described in the Materials and Methods section. We defined this value as the amplitude index, and the mean amplitude index in 10 wild type ovules was 2076.8 ± 730.1 during Phase 1-2 (Supplementary Fig. S6). Quantification of the amplitude index across 10 replicates for each mutant revealed a significant reduction compared with the wild type (*aru-1*, 699.3 ± 250.5; *alg10-1*, 941.1 ± 406.5) similar to those in *fer-4* (574.8 ± 81.8) (Supplementary Fig. S6). However, in many ovules of both mutants, elevated cytosolic calcium levels in synergid cells prior to pollen tube rupture (Phase 2) were maintained.

To investigate the cause of attenuated calcium oscillations, we examined the localization of proteins involved in pollen tube reception. Marker lines for LORELEI, NORTIA, and ANJEA were generated and introduced into *aru-1* or *alg10-1* mutants by crossing. LORELEI and ANJEA contain putative glycosylation sites, and NORTIA is known to traffic to the synergid cell plasma membrane via the N-glycosylated receptor FERONIA (Ju et al. 2021). Consistent with Müller et al. (2016), the localization patterns of LORELEI and NORTIA appeared normal in *aru-1*, and similar results were observed in *alg10-1* (Supplementary Fig. S6A, C). ANJEA was also localized to the filiform apparatus in both mutants, similar to that in the wild type (Supplementary Fig. S6B).

When we performed aniline blue staining of ovules, many mature ovules exhibited increased callose deposition at the filiform apparatus in both mutants compared to the wild type (Fig. 3A, C). Glutaraldehyde fixation further showed that the filiform apparatus was thicker than that in the wild type (Fig. 3B). This phenotype resembles that of *evan* and *turan*, mutants defective in upstream steps of the N-linked glycosylation pathway (Lindner et al. 2015). Increased callose accumulation was also observed in approximately 50% of *alg10-1* heterozygous ovules (Fig. 3C), suggesting that this phenotype is caused by defects in the female gametophyte. During Phase 1, pollen tubes directly contact the filiform apparatus. Therefore, increased callose deposition and structural alterations of the filiform apparatus could affect communication between the pollen tube and the synergid cell required for normal calcium oscillations.

**Fig. 3.**
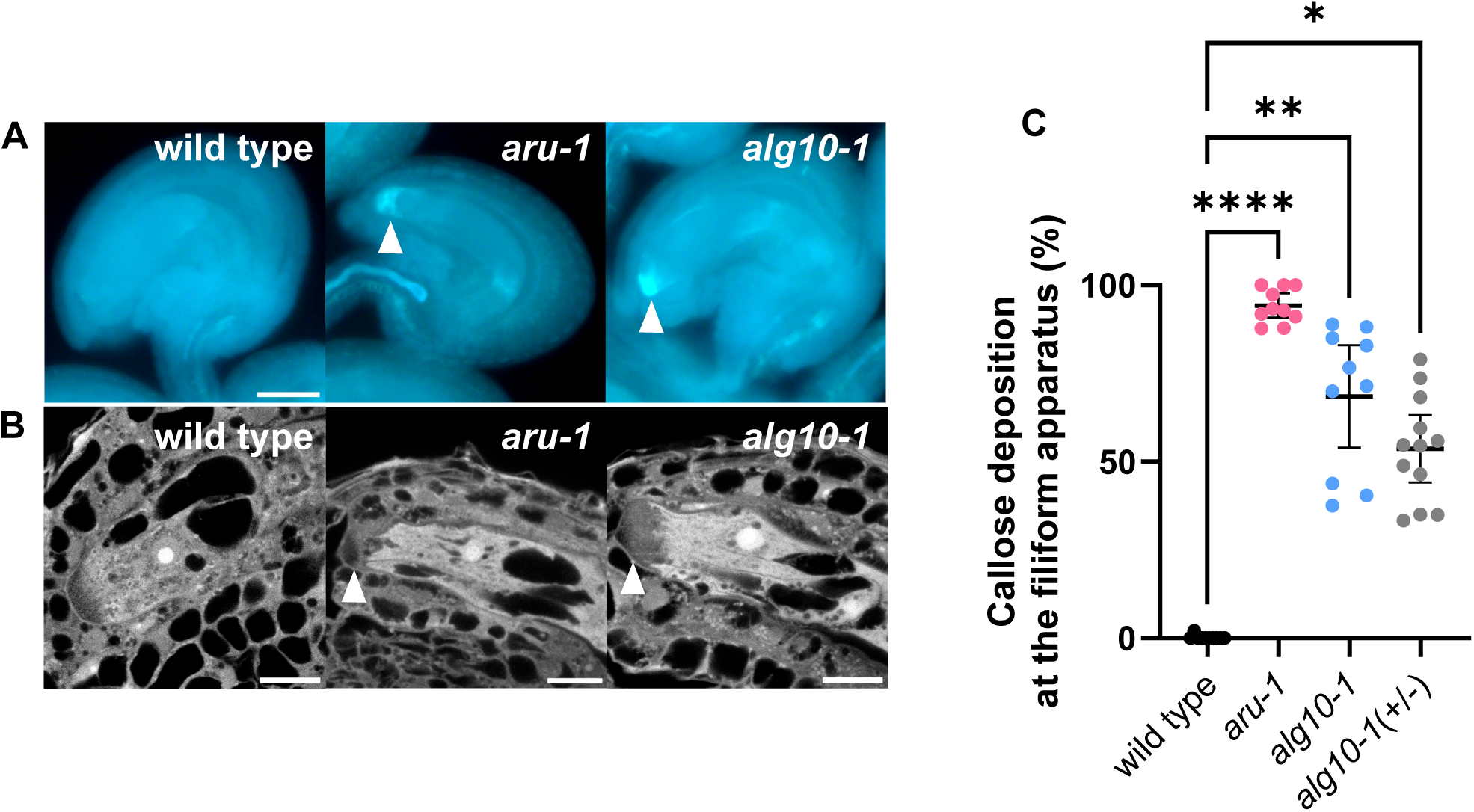
*aru-1* and *alg10-1* ovules display increased callose accumulation at the filiform apparatus caused by abnormalities in the female gametophyte. (A) Aniline blue staining of callose in mature ovules. Arrowheads indicate aberrant callose deposition at the filiform apparatus. Scale bar, 20 μm. Note that the vascular bundles within the funiculus were also stained. (B) Confocal images of the filiform apparatus of glutaraldehyde-fixed mature ovules. Arrowheads indicate abnormal structures at the filiform apparatus. Scale bar, 10 μm. (C) Frequency of ovules displaying increased callose deposition at the filiform apparatus in wild-type control (n = 10), *aru-1* (n = 10), *alg10-1*−/− (n = 10), and *alg10-1*+/− (n = 12). Data show mean ± 95% CI. Each point represents an individual pistil. Statistical significance was determined using the Kruskal–Wallis test, followed by Dunnett’s multiple comparisons test. ****P < 0.0001, **P = 0.0013, *P = 0.0268. Note that for the *alg10-1* homozygous mutant, the count did not include ovules exhibiting aberrant callose accumulation within the embryo sac, as shown in Supplementary Fig. S8.

### aru-1 does not show reduced fertility, and the reduced fertility of alg10-1 is caused by female sporophytic mutation

To date, all mutants exhibiting defects in calcium oscillations in synergid cells have shown reduced fertility (Ngo et al. 2014, Desnoyer et al. 2024, Gao et al. 2022, Ponvert and Johnson 2024). To elucidate the precise role of synergid cell calcium oscillations during pollen tube reception, we examined the fertility phenotypes of the *aru-1* and *alg10-1* mutants. Notably, consistent with previous observations (Müller et al. 2016), the seed set of *aru-1* was not reduced compared with that of the wild type (Fig. 4A, B). In contrast, fruits of *alg10-1* were shorter and wider than those of the wild type, accompanied by an approximately 20% reduction in seed set (Fig. 4A, B). As reported in a previous study (Farid et al. 2011), *alg10-1* exhibits a semi-dwarf phenotype; reduced ovule number was also observed (Fig. 4A). These phenotypes were not observed in the *aru-1*. To determine whether the fertility defect of *alg10-1* originated from the male or female parent, we performed reciprocal crosses. When homozygous *alg10-1* plants were used as the female parent, seed set was significantly reduced (Fig. 4C). In contrast, when *alg10-1* was used as the pollen donor, no reduction in seed set was observed. To further determine whether the female-specific defect in *alg10-1* arises from sporophytic tissues or the gametophyte, we examined seed set in *alg10-1* heterozygotes and assessed transmission efficiency. The *alg10-1* heterozygote showed no reduction in fertility in self-pollination or when pollinated with wild-type pollen. Consistent with these results, 57.6% (87/151) of the seeds obtained by pollinating pistils of the *alg10-1* heterozygote with wild-type pollen inherited the mutant allele.

**Fig. 4.**
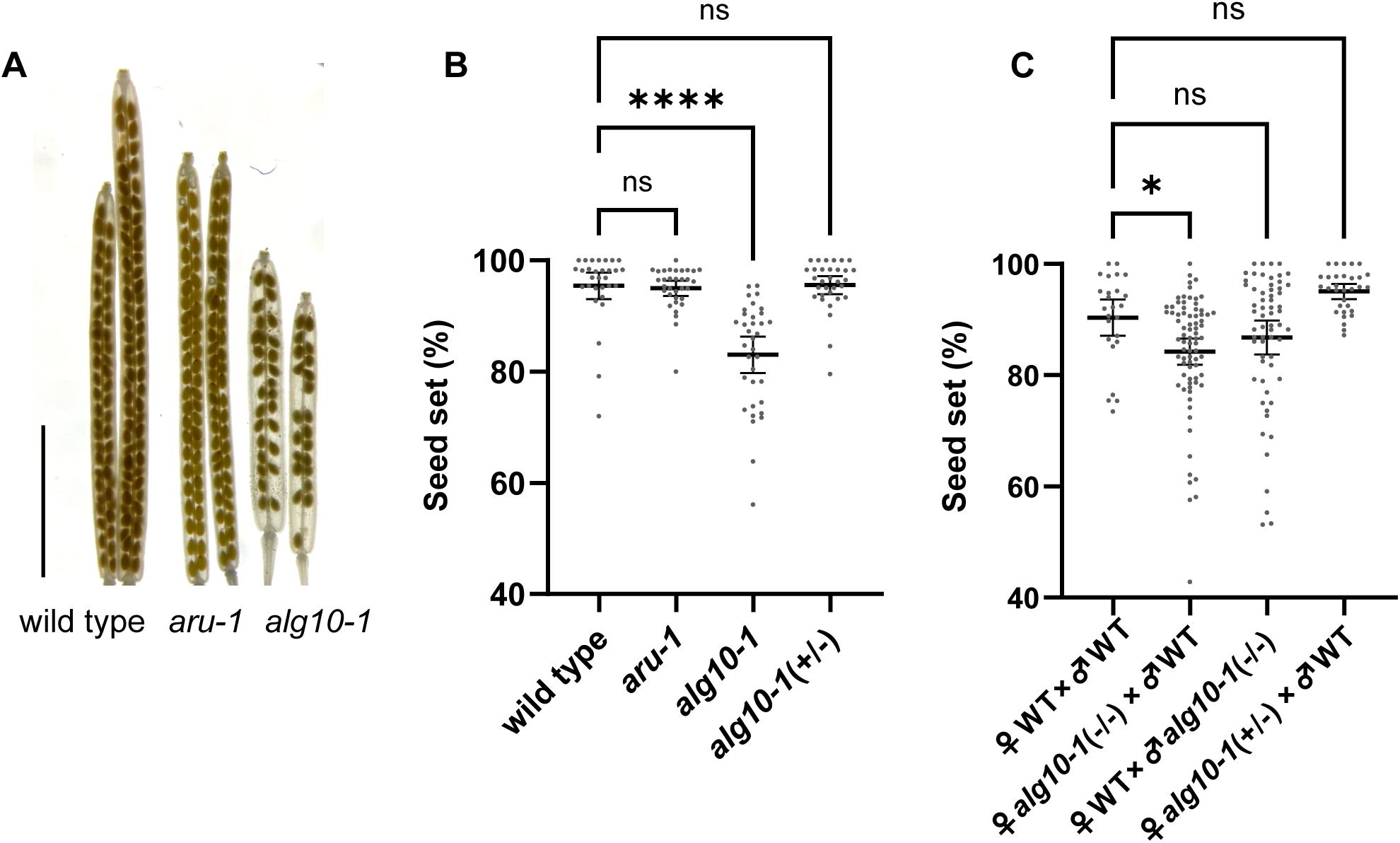
*aru-1* does not show reduced fertility and the reduced fertility of *alg10-1* is not caused by abnormalities in the female gametophyte. (A) Fruits from *aru-1*, *alg10-1,* and the wild-type control. Scale bar, 5 mm. (B) Seed set of self-pollinated wild type (n = 30), *aru-1* (n = 33), alg*10-1* (n = 35), and *alg10-1* (+/−) (n = 32). (C) Seed set of fruits obtained from reciprocal crosses. From the experimental group on the left: n = 25, n = 78, n = 62, and n = 31, respectively. (B-C) Data show mean ± 95% CI. Each point represents an individual fruit. Statistical significance was determined using the Kruskal–Wallis test, followed by Dunnett’s multiple comparisons test. ****P < 0.0001, *P = 0.013, ns = not significant.

To further investigate female defects in *alg10-1*, we examined pollen tube guidance efficiency and ovule morphology using aniline blue staining (Supplementary Fig. S8). One day after pollination, 6.5 ± 5.2% of *alg10-1* ovules failed to attract pollen tubes (n = 15 pistils), which was higher than that in wild-type ovules (2.2 ± 2.9%). In *alg10-1* ovules, aberrant callose accumulation due to a defect in embryo sac formation was observed (6.7 ± 9.3%; n=10), indicating that reduced fertility in *alg10-1* should be a developmental ovule defect (Supplementary Fig. S8).

### alg10-1 exhibits polytubey caused by defects in female sporophytic tissues

Furthermore, aniline blue staining revealed that 42.1 ± 9.2% of *alg10-1* ovules were targeted by more than two pollen tubes (polytubey) at 24 hours after pollination (HAP) with wild-type pollen (n = 15), which was significantly higher than that with wild-type ovules (Fig. 5A–C). Because polytubey was observed as early as 6 HAP, this defect is unlikely to be explained solely by recovery following failed acceptance of the first pollen tube (Kasahara et al. 2012) (Fig. 5B). While many polytubey cases of *fer-4* or *lre-5* were accompanied by overgrowth of the first pollen tube, no increase in the overgrowth rate was observed in *alg10-1* (consistent with Zupunski, 2020). To determine whether the elevated polytubey rate in *alg10-1* pistils is caused by defects in sporophytic tissues or the gametophyte, we examined polytubey in *alg10-1* heterozygotes. The *alg10-1* heterozygote did not exhibit a polytubey phenotype when pollinated with wild-type pollen (Fig. 5C), suggesting that the polytubey is due to a sporophytic defect.

**Fig. 5.**
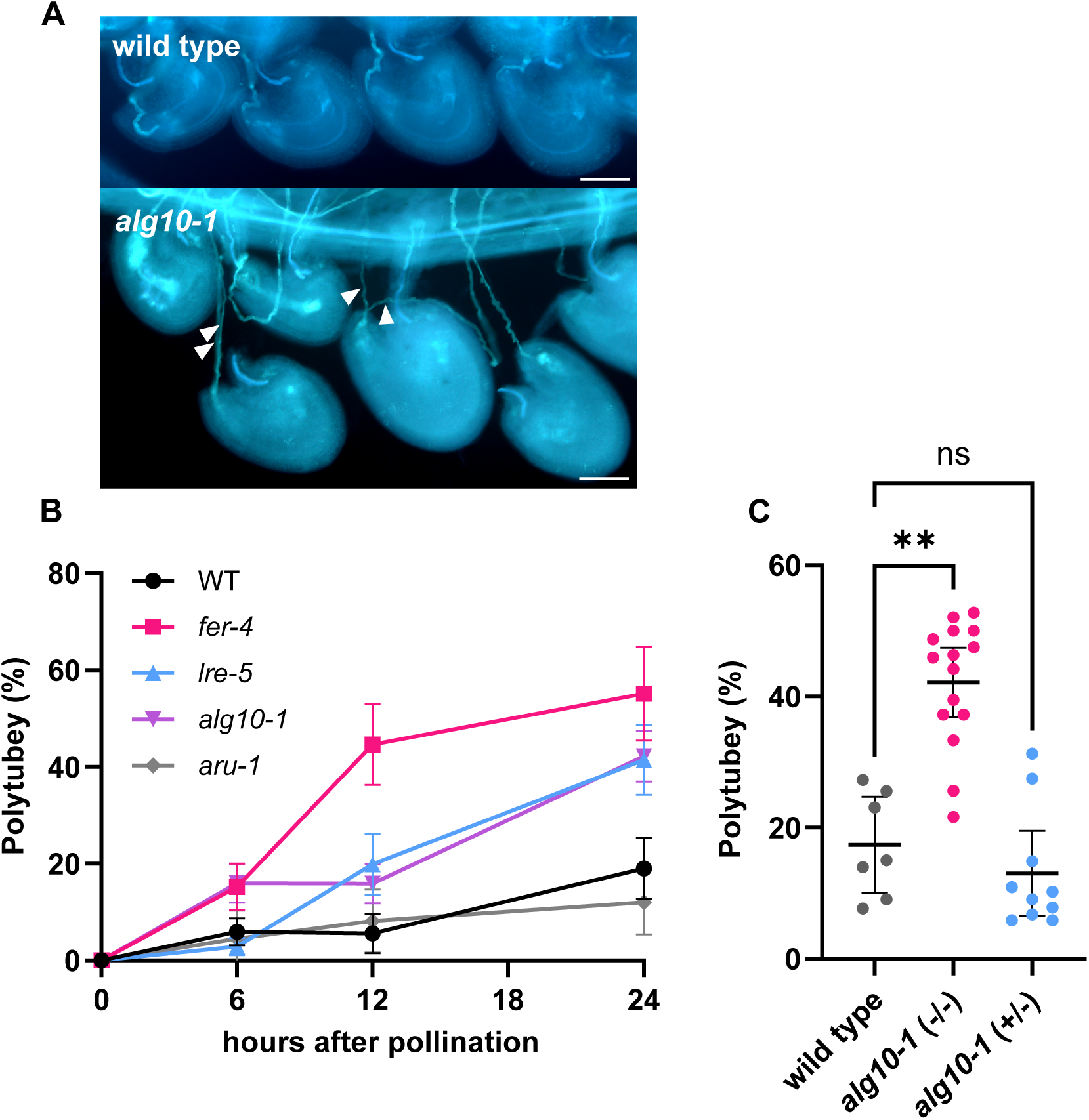
*alg10-1* exhibits early-onset polytubey caused by female sporophytic tissue. (A) Aniline blue staining of multiple pollen tubes in *alg10-1* and wild-type control. The females were pollinated with wild-type pollen. Arrowheads indicate pollen tubes targeting a single ovule. Scale bar, 50 μm. (B) Polytubey rate (%) in wild type, *fer-4*, *lre-5*, *aru-1,* and *alg10-1* at 6, 12, and 24 h after pollination. “n” refers to the number of pistils. Data are mean ± 95% CI. (C) Frequency of ovules with multiple pollen tubes entering homozygous (−/−) or heterozygous (+/−) *alg10-1* mutants. Data are mean ± 95% CI. Each point represents an individual pistil. Statistical significance was determined using the Kruskal–Wallis test, followed by Dunnett’s multiple comparisons test. **P = 0.004, ns = not significant.

### Sporophytic tissues of the ovule exhibit characteristic calcium signaling during pollen tube reception

As suggested by live-cell imaging, ovular sporophytic tissues contacting an attracted pollen tube can potentially contribute to a polytubey block even before pollen tube reception by the synergid cell (Mizuta et al. 2024). We investigated whether calcium signaling in sporophytic tissues is involved in pollen tube reception by the ovule. To monitor calcium dynamics throughout the ovule, GCaMP6s was expressed under the control of the ubiquitously expressed *Ribosomal Protein Subunit 5A* (*RPS5A*) promoter (Adachi et al. 2011). The ovule consists of a female gametophyte (n), containing two synergid cells, an egg cell, and a central cell, surrounded by diploid sporophytic tissues (2n) comprising the outer and inner integuments and the nucellus. To distinguish signals from gametophyte and sporophytic tissues, we used plants homozygous or heterozygous for *pRPS5A::GCaMP6s*. Homozygous ovules expressed GCaMP6s in all cells, whereas approximately half of the ovules from heterozygous plants expressed GCaMP6s exclusively in sporophytic tissues. Thus, a comparison of these genotypes enabled the discrimination of sporophyte-derived signals. In addition, Z-stack imaging allowed the separation of tissue-specific signals based on focal planes.

Using these approaches in a semi-*in vivo* assay, we found that ovules exhibited several characteristic calcium waves within sporophytic tissues. Prior to pollen tube arrival, calcium waves occurred spontaneously and sporadically in sporophytic tissues (Fig. 6A). Upon pollen tube arrival at the micropyle, a calcium wave was initiated in the micropylar region and propagated toward the chalazal region, reaching the funiculus (Fig. 6B). Following pollen tube rupture, a calcium wave was initiated near the synergid cells and propagated toward the chalazal end (Fig. 6C). This wave, the pollen tube discharge-triggered calcium wave, was also observed in ovules from heterozygous plants expressing GCaMP6s exclusively in sporophytic tissues. This calcium wave was not observed in cases when pollen tube rupture failed and overgrowth occurred.

**Fig. 6.**
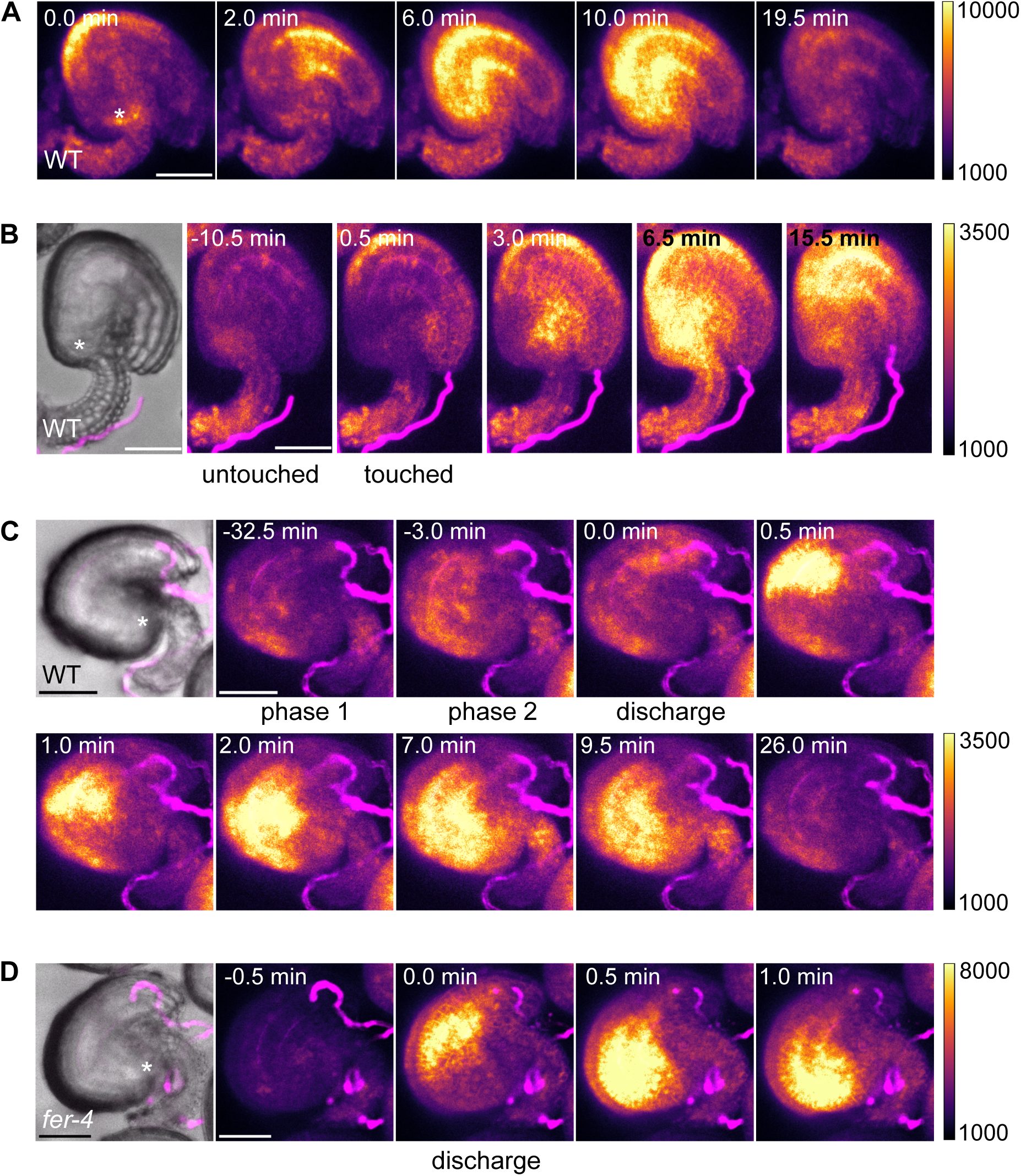
Sporophyte tissues of the ovule exhibit characteristic calcium waves corresponding to pollen tube dynamics. (A) A time-lapse series showing a calcium wave in the default mode. An ovule expressing GCaMP6s is visible. Time point 0 is the start of the observation. Scale bar, 50 μm. (B) A time-lapse series showing a calcium wave triggered by the arrival of the pollen tube. (C) A time-lapse series showing a calcium wave triggered by sperm cell discharge. (D) A time-lapse series showing a calcium wave triggered by sperm cell discharge in *fer-4* ovule. (B-D) An ovule expressing GCaMP6s and a pollen tube expressing mApple (magenta) are shown. The left panel shows a merged bright-field and confocal image of an ovule and pollen tube (magenta). Time point 0 indicates when the pollen tube ruptured. Scale bar, 20 μm. (C-D) Asterisks indicate the chalazal region of the ovule.

### FERONIA is not required for calcium waves associated with pollen tube rupture

FERONIA plays a crucial role in pollen tube reception by synergid cells (Huck et al. 2003, Rotman et al. 2003, Escobar-Restrepo et al. 2007). It is also expressed in the septum and sporophytic tissues of the ovule, where it contributes to the polytubey block (Duan et al. 2020, Zhong et al. 2022). To investigate the role of FERONIA in calcium waves within sporophytic tissues of the ovule, we introduced *pRPS5A::GCaMP6s* into the *fer-4* mutant and performed a semi-*in vivo* assay. In *fer-4*, irregular spontaneous calcium waves prior to pollen tube arrival and rapid calcium waves upon pollen tube rupture were observed under semi-*in vivo* conditions, similar to those in the wild type (Fig. 6D). In contrast, calcium waves triggered by the pollen tube arrival were not clearly detected.

## Discussion

In this study, we visualized calcium dynamics in the ovule under semi-*in vivo* conditions using GCaMP6s expressed either ubiquitously or specifically in synergid cells. This approach enabled us to monitor the transition from pollen tube arrival to sperm release. We identified mutants that exhibited attenuated calcium oscillations in synergid cells and demonstrated that such attenuation does not necessarily impair fertilization. In addition, we identified multiple classes of calcium waves in sporophytic tissues of the ovule and showed that some of these were independent of FERONIA. Together, these findings suggest that ovular calcium signaling during pollen tube reception is multi-layered, involving both gametophytic and sporophytic components.

### N-linked glycosylation is required for calcium oscillations in synergid cells triggered by pollen tube arrival at the micropyle

We found that loss-of-function mutants of the N-linked glycosylation-related OST3/6 subunit (*aru-1*) and ALG10 (*alg10-1*) exhibited attenuated calcium oscillations in synergid cells during pollen tube reception under semi-*in vivo* conditions. These observations suggest that the N-linked glycosylation machinery is required for the generation of proper calcium oscillations in synergid cells. As described above, N-linked glycosylation affects the folding, stability, transport, and activity of many cell surface and secreted proteins (Helenius and Aebi 2004, Sato et al. 2017). For example, previous studies in *Arabidopsis thaliana* have shown that a loss-of-function mutant of ALG3, which functions upstream of ALG10 and OST in N-glycan synthesis, causes defects in the glycosylation of immune receptors on the cell surface (Trempe et al. 2016). This, in turn, results in the absence of the calcium increase that normally occurs upon recognition of pathogen-derived molecules. The reduced calcium oscillations observed in synergid cells in this study may similarly result from impaired glycosylation of cell surface receptors, leading to weaker perception of pollen tube-derived signals. In this study, the localization of LORELEI, ANJEA, and NORTIA, which are involved in pollen tube reception, was not altered in synergid cells of *aru-1* and *alg10-1*. Previous studies have also reported that the localization and expression levels of FERONIA remain unchanged in *aru-1* (Müller et al. 2016). However, as suggested previously (Müller et al. 2016), the absence of N-linked glycosylation does not necessarily affect protein localization and may instead impair protein function or stability.

Furthermore, increased callose accumulation at the filiform apparatus was observed in many ovules of *aru-1* and *alg10-1*, and the filiform apparatus appeared thicker than in the wild type. The filiform apparatus functions as a communication interface between synergid cells and the pollen tube. These morphological abnormalities may affect the organization of the cell membrane and surface receptors and may impair communication with the pollen tube required to trigger an increase in calcium levels in synergid cells.

### The reduction in amplitude of calcium oscillations in synergid cells during pollen tube reception does not reduce fertility

Calcium oscillations in synergid cells are considered a hallmark of pollen tube reception. However, *aru-1* showed no significant reduction in fertility compared with the wild type. In *alg10-1*, fertility decreased by approximately 20%, although this appeared to be mainly attributable to defects in female sporophytic tissues. These data suggest that calcium oscillations and their precise patterns in Phase 1 may not serve as a strict trigger for pollen tube rupture but rather as a consequence of complex male–female molecular interactions. In contrast, the sustained elevation of calcium levels prior to pollen tube–synergid cell rupture was largely maintained in *aru-1* or *alg10-1* synergid cells. Ngo et al. (2014) reported that reducing calcium levels in synergid cells using the chelator BAPTA-AM caused pollen tube overgrowth, suggesting that an increase in calcium levels in synergid cells is required for pollen tube rupture. Our results further suggest that elevated calcium levels (Phase 2), rather than the oscillatory pattern itself, are critical for pollen tube rupture.

Meanwhile, previous studies have reported that pollination with pollen derived from the closely related species *Arabidopsis lyrata* results in a reduced frequency of successful pollen tube reception and an increased incidence of pollen tube overgrowth in *aru-1* and *alg10-1*, and that this overgrowth phenotype is attributable to synergid cells (Müller et al. 2016, Zupunski, 2020). These findings suggest that N-linked glycosylation in synergid cells contributes to promote the rupture of heterospecific pollen tubes. Recently, Ponvert and Johnson (2024) analyzed the frequency and amplitude of calcium oscillations in synergid cells and reported that the oscillatory pattern remains largely unchanged upon the reception of heterospecific pollen tubes. This is consistent with our observation that Phase 2 is more critical, and the pattern of calcium oscillation in Phase 1 is not a constraint of Phase 2.

### N-linked glycosylation in female sporophytic tissues is involved in the polytubey block

Our results further reveal that *alg10-1* exhibits a high incidence of early-onset polytubey. As this phenotype appears to originate in female sporophytic tissues, these findings suggest that N-linked glycosylation in the female sporophyte is required for establishing the polytubey block, independently of synergid calcium oscillations. The polytubey block is a complex, multi-layered signaling process (reviewed in Sugi and Maruyama 2023). As described above, FERONIA expressed in the septum blocks additional pollen tube emergence (Zhong et al. 2022), and FERONIA expressed in synergid cells also suppresses polytubey (Duan et al. 2020). Recent studies have further shown that after the first pollen tube reaches the synergid cell or micropyle, the release of pollen tube attractants derived from synergid cells, including LURE peptides, is either halted or the attractants are actively degraded, thereby preventing the attraction of subsequent pollen tubes (Yu et al. 2021, Duan et al. 2020, Maruyama et al. 2015, Völz et al. 2013). In addition, mutants that frequently exhibit polytubey include double mutants of ANJEA and HERK1, which are involved in the perception of pollen tube-derived RALF peptides (Galindo-Trigo et al. 2020, Zhong et al. 2022), as well as mutants of the glycoprotein AGP4 (also known as JAGGER) (Pereira et al. 2016). Notably, FERONIA, ANJEA, and HERK1 contain putative N-linked glycosylation sites, raising the possibility that impaired glycosylation of these proteins contributes to the polytubey phenotype observed in *alg10-1*. In contrast, *alg10-1* does not exhibit significant pollen tube overgrowth; thus, its phenotype differs from that of the loss-of-function mutants of these receptors.

### Sporophytic tissues of the ovule sense contact and rupture of the pollen tube and propagate signals toward the funiculus

Under semi-*in vivo* conditions, we observed calcium waves in sporophytic tissues of the ovule elicited upon contact and rupture of the pollen tube. These calcium waves propagate over long distances, including along the funiculus. In addition, our observation that certain calcium waves persist in the *fer-4* mutant indicates that not all calcium signaling events during pollen tube reception depend on FERONIA. Furthermore, since the amplitude of calcium oscillations in the synergid cells of *fer-4* is reduced, there does not appear to be a simple correlation between the calcium oscillation patterns in the synergid cells and the calcium waves in the sporophyte tissues of the ovule. Previous studies on the polytubey block have shown that the signal indicating successful reception of the first pollen tube is transmitted to the funiculus and septum, thereby inhibiting the growth of additional pollen tubes toward the same ovule (Mizuta et al. 2024). However, the mechanism underlying this long-distance signaling remains unclear. This signaling may be mediated by diffusible molecules, such as nitric oxide (NO) (Duan et al. 2020), or by cell–cell interactions within sporophytic tissues of the ovule. Calcium ions function as pivotal second messengers and are involved in electrical signal propagation in plants (Vodeneev et al. 2016); thus, the observed calcium waves may reflect intercellular communication within the ovule. These calcium waves may trigger downstream responses, including the polytubey block, and could represent a form of long-distance signaling mediated through sporophytic tissues. These observations suggest that calcium dynamics visualize previously uncharacterized intercellular communication events involved in pollen tube reception and subsequent responses. Further analysis by manipulating calcium dynamics in the ovule would provide insights for elucidating the mechanisms that ensure precise one-to-one pollen tube reception by the ovule.

## Materials and Methods

### Plant materials

*Arabidopsis thaliana* strain Columbia (Col-0) was used as a wild-type plant. The seeds for *alg10-1* (SAIL_515_F10) and *fer-4* (GABI-GK106A06) were obtained from the *Arabidopsis* Biological Resource Center at Ohio State University (Columbus, OH, USA), and *aru-1* (SALK_067271) was obtained from The Nottingham Arabidopsis Stock Centre at the University of Nottingham (Loughborough, UK). *pLAT52::mApple* (Kurihara et al. 2015) has been previously described. Col-0 was transformed with *pMYB98::GCaMP6s*, *pMYB98::LORELEIΔω: mClover*, *pMYB98::ANJEA:mRuby3,* and *pMYB98::NORTIA:mRuby3* using *Agrobacterium*-mediated transformation, as described previously (Narusaka et al. 2010). Double mutants/marker lines were generated by crossing and confirmed by PCR genotyping or microscopy-based phenotyping.

### Growth conditions

Seeds were sterilized with 70% (v/v) ethanol for 10 min and sown on agar plates containing 1/2 MS medium (1/2 MS, 0.8% [w/v] agar, and 1% [w/v] sucrose, pH adjusted to 5.8 with KOH). Plates were incubated at 4 ℃ for 1–2 days and then transferred to a growth chamber at 22 ℃ under 24 h of light. Ten-day-old seedlings were transferred into a mixture of vermiculite and potting compost in a growth room with LED light under long-day condition (16 h light/8 h dark cycle) at 21 ℃.

### Plasmid construction

To generate the translational reporters for *LORELEI*, *NORTIA,* and *ANJEA* genes, genomic or cDNA sequences of each gene were cloned into pPZP211 (Hajdukiewicz et al. 1994) or pMDC99 (Curtis and Grossniklaus 2003). The GCaMP6s coding sequence was also cloned into pPZP211 vector with the promotor regions of *MYB98* (Kasahara et al. 2005) or *RPS5A* (Adachi et al. 2011).

### Ca^2+^ imaging with semi-in vivo assay

A semi-*in vivo* assay was performed as described by Hamamura et al. (2014), with some modifications. Stage 12 flowers were emasculated 17–20 h before dissection. 70 μL of pollen germination media (5 mM KCl, 0.01% H_3_BO_3_, 5 mM CaCl_2_, 1 mM MgSO_4_, 10% Sucrose, 1.5% NuSeive GTG Agarose, pH 7.6) (Boavida and McCormick 2007) was poured in the well (1 cm Length x 1.5 cm Width x 0.1 cm Hight) on 25×50 mm cover glass (C025501, Matsunami) and allowed to solidify. The well was covered with another cover glass to maintain humidity. The excised pistils were pollinated under a stereomicroscope and cut with a 26G needle at the junction between the style and ovary. Excised stigma was placed on the medium immediately. Ovules with their funiculus were removed from septum and immediately put on the medium within 150 μm from the pistil excision site. Synergid calcium oscillations were observed 2.5–4 h after pollination at 21°C in the dark. We usually started time-lapse imaging just before the pollen tubes reached the ovules. Synergid cells and pollen tubes were imaged using an inverted microscope (IX-83; Olympus) equipped with an automatically programmable XY stage (BioPrecision; Ludl Electronic Products Ltd, Hawthorne, NY), a spinning-disk confocal system (CSU-W1; Yokogawa Electric), 488 nm and 561 nm (Sapphire; Coherent) LD lasers, and an EM-CCD camera (ImagEM 1K; Hamamatsu Photonics, Shizuoka, Japan). Images were taken with a 30 s interval and continuing for 6 h using multiple z-planes (5 μm intervals) and 13 planes with a ×20 objective lens (UPlanSApo 20x/0.75). Exposure times were 90 ms for GCaMP6s and 70 ms for mApple. The imaging system was controlled by MetaMorph software version 7.8.10.0 (Universal Imaging Corp., Downingtown, PA).

### Image processing

The fluorescence intensity of GCaMP6s in the synergid cells was measured using the following procedure. Hyperstack images were created using the Fiji software (Schindelin et al. 2012) and subsequently projected using the Sum slices method. The region of the synergid cells was identified using Otsu thresholding and a median filter, which were then applied to the sum projection image to measure the fluorescence intensity within the region. The fluorescence intensity of GCaMP6s was calculated by subtracting the background brightness from the obtained values. The distance by which pollen tubes elongated from the entrance of the micropyle was calculated by manually tracking pollen tube tips on z-projected images using the MTrackJ plugin in the Fiji software. All videos and images were generated using Fiji.

### Amplitude index

To quantitatively compare the amplitude of calcium oscillations in synergid cells in Phase 1 and 2, we measured the “amplitude index” using the following procedure. Using the GCaMP6s fluorescence intensity of the synergid cells obtained by the method described above, we calculated the average change in fluorescence intensity at each time point from the arrival of the pollen tube at the filiform apparatus until the rupture of the receptive synergid cell. Specifically, we obtained the absolute value of the difference in the GCaMP6s fluorescence intensity between each time point and the time point 30 s prior, and then divided the sum of these values by the number of time points.

### Localization analysis using confocal microscopy

Stage 12–13 flowers were emasculated 17–20 h before observation. The following day, ovules were collected and mounted on slides in 1/2 MS medium containing 5% sucrose. Images were captured using a Leica STELLARIS 8 confocal laser scanning microscope with filters for ALEXA 488 or TagRFP.

### Aniline blue staining

Stage 12–13 flowers were emasculated 17–20 h before pollination. Pistils were pollinated with wild-type pollen. At certain hours after pollination, pistils were collected and fixed in 9:1 ethanol:acidic acid overnight. They were rinsed three times with dH_2_O and then softened with 8 M NaOH overnight at 4 ℃. Pistils were washed with dH_2_O three times and stained with a decolorized aniline blue solution (0.1% aniline blue, 0.1 M K_3_PO_4_) for more than 1 h in the dark. Before microscopic observation, an ovary wall was removed with a 26G needle under a stereomicroscope and the samples were mounted with a decolorized aniline blue solution. The samples were analyzed using a Zeiss AxioObserver Z1 inverted microscope equipped with an ultraviolet filter set (Filter Set 02).

### Glutaraldehyde fixation

To observe the inner structure of the ovules, glutaraldehyde fixation was conducted as previously described (Sandaklie-Nikolova et al. 2007, Jia et al. 2020) with some modifications. We harvested the pistils on double-sided tape on a slide glass and opened the ovary walls by making cuts on both sides of the pistil replum using a surgical needle to expose the ovules to the fixative. They were immediately soaked in the fixation solution (4% glutaraldehyde in 12.5 mM cacodylate buffer, pH 6.9) and incubated overnight at room temperature. After fixation, they were dehydrated sequentially with 10%, 30%, 50%, 70%, 90% and 100% ethanol, for 20 min at each concentration, and then cleared in 2:1 (v/v) benzyl benzoate: benzyl alcohol for 1 h. Ovules were then dissected out using a 26G needle, mounted in immersion oil, and observed under a Leica STELLARIS 8 confocal laser scanning microscope with filters for ALEXA 488, 488 nm excitation and a 515- to 565-nm emission filter.

### Seed set analysis

Siliques were cleared as previously described (Flores-Tornero et al. 2021). Briefly, almost matured siliques were collected in a fixative solution (absolute ethanol:glacial acetic acid (3:1)) and left overnight at 4°C. Fixed siliques were washed with 90% ethanol for 10 min and replaced with 70% ethanol. Samples were stored at 4°C until observation. number of seeds was counted under a stereomicroscope (SZX-ZB7; EVIDENT) equipped with a light source from the bottom.

### Transmission efficiency

To detect abnormalities in the female gametophyte, the transmission efficiency was calculated using the following method. Flowers of *alg10-1* (+/−) at stage 12-13 were emasculated. The following day, the pistils were hand-pollinated with wild-type pollen. Approximately two weeks later, seeds were collected from mature fruits and sown on 1/2 MS plates, as described above. The seedlings were genotyped by PCR, and the proportion of plants that inherited the mutant allele was calculated.

### Statistical analysis

Data visualization and statistical analyses were performed using GraphPad Prism v11.0.0 (GraphPad Software, San Diego, CA, USA). Statistical significance between the two groups was determined using the Mann-Whitney test. To determine significant differences between more than three groups, the Kruskal–Wallis test followed by Dunnett’s multiple comparison test was employed in this study.

## Acknowledgements

We thank Tadasuke Chiba (EVIDENT) and Yoshitaka Sekizawa (Yokogawa Electric Corporation) for their technical advice regarding confocal microscopy.

