## Supplemental Figures for "Calcium Dynamics During Pollen Tube Reception in *Arabidopsis* Ovules"

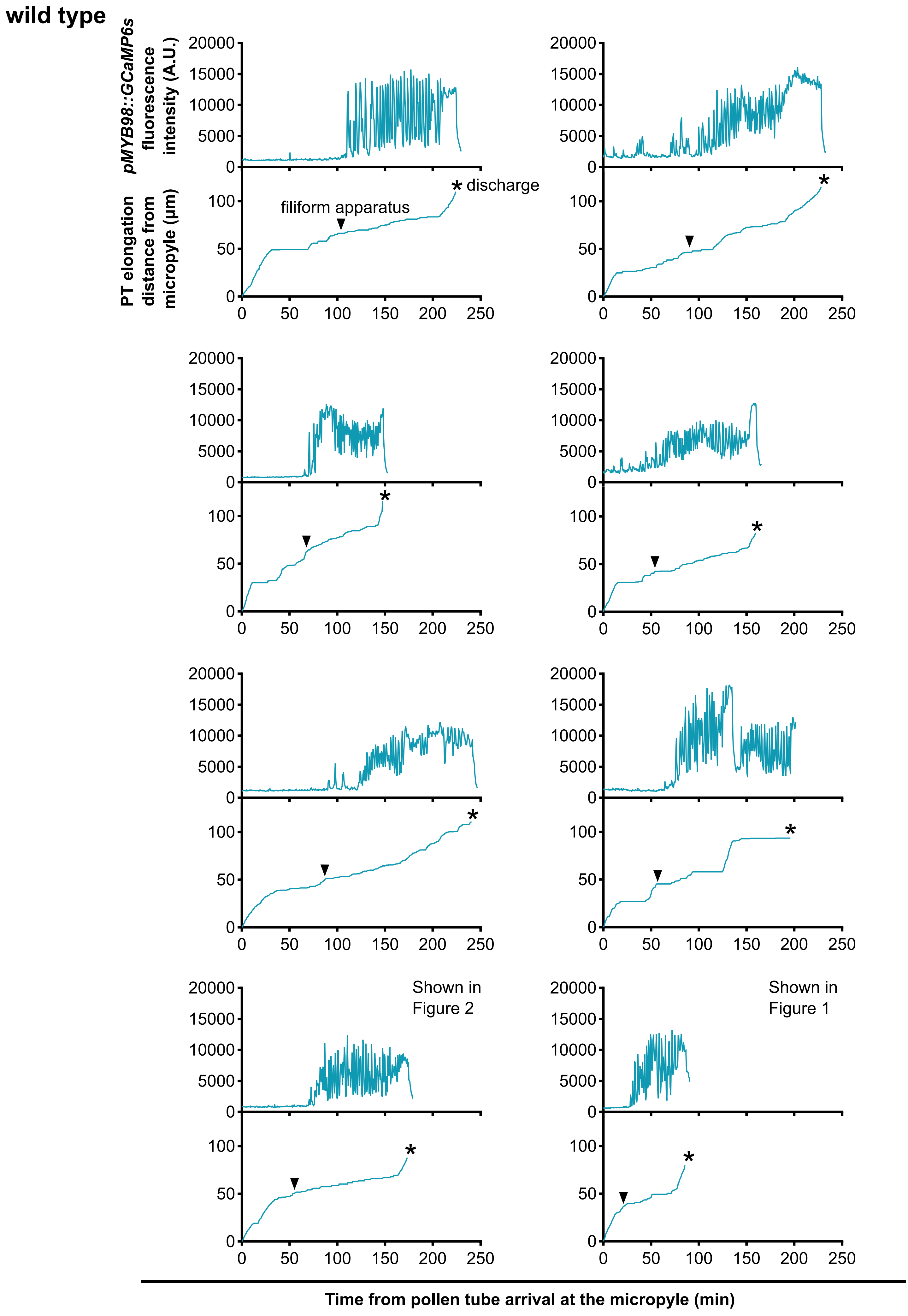


(Continued on next page)

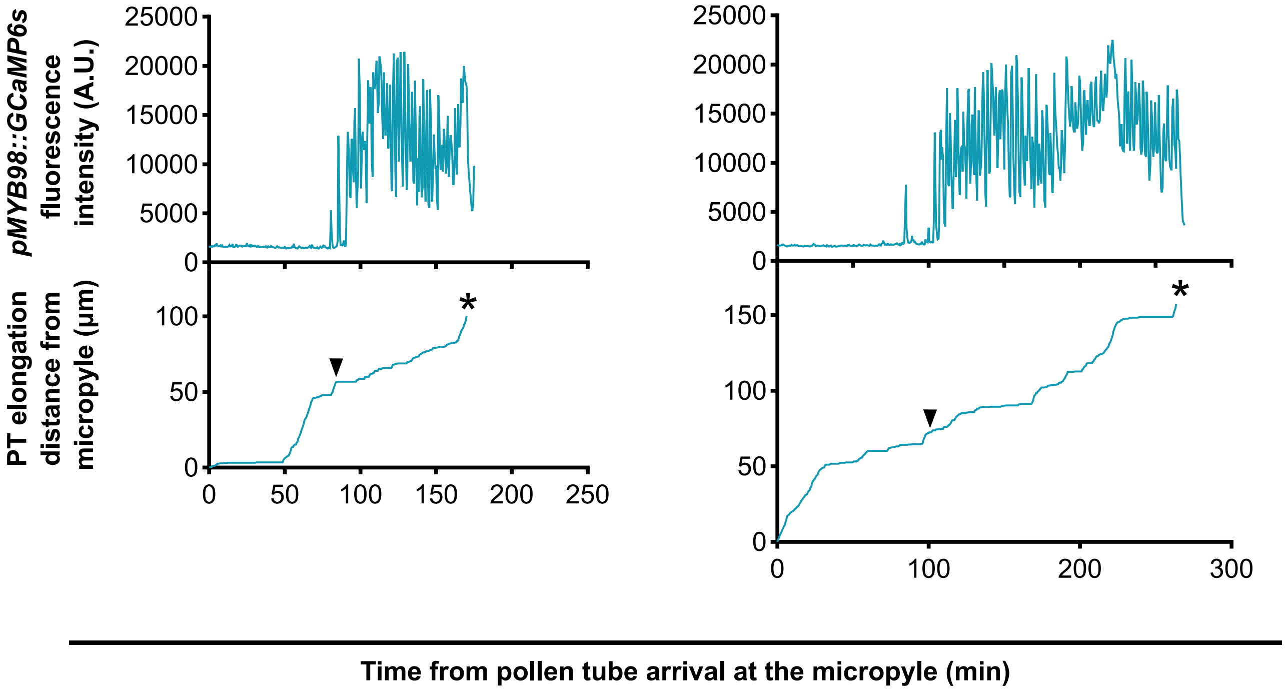


**Supplementary Figure S1** Ten replicate time-lapse calcium ion traces of *pMYB98:GCaMP6s* in the wild type. The upper graphs show the fluorescence intensity of GCaMP6s in the synergid cells, and the lower graphs show the pollen tube elongation distance from the micropyle. All were pollinated with wild-type pollen. The arrowheads labeled "filiform apparatus" indicate when the pollen tube tip reached the filiform apparatus. Astarisks indicate pollen tube-synergid cell rupture.

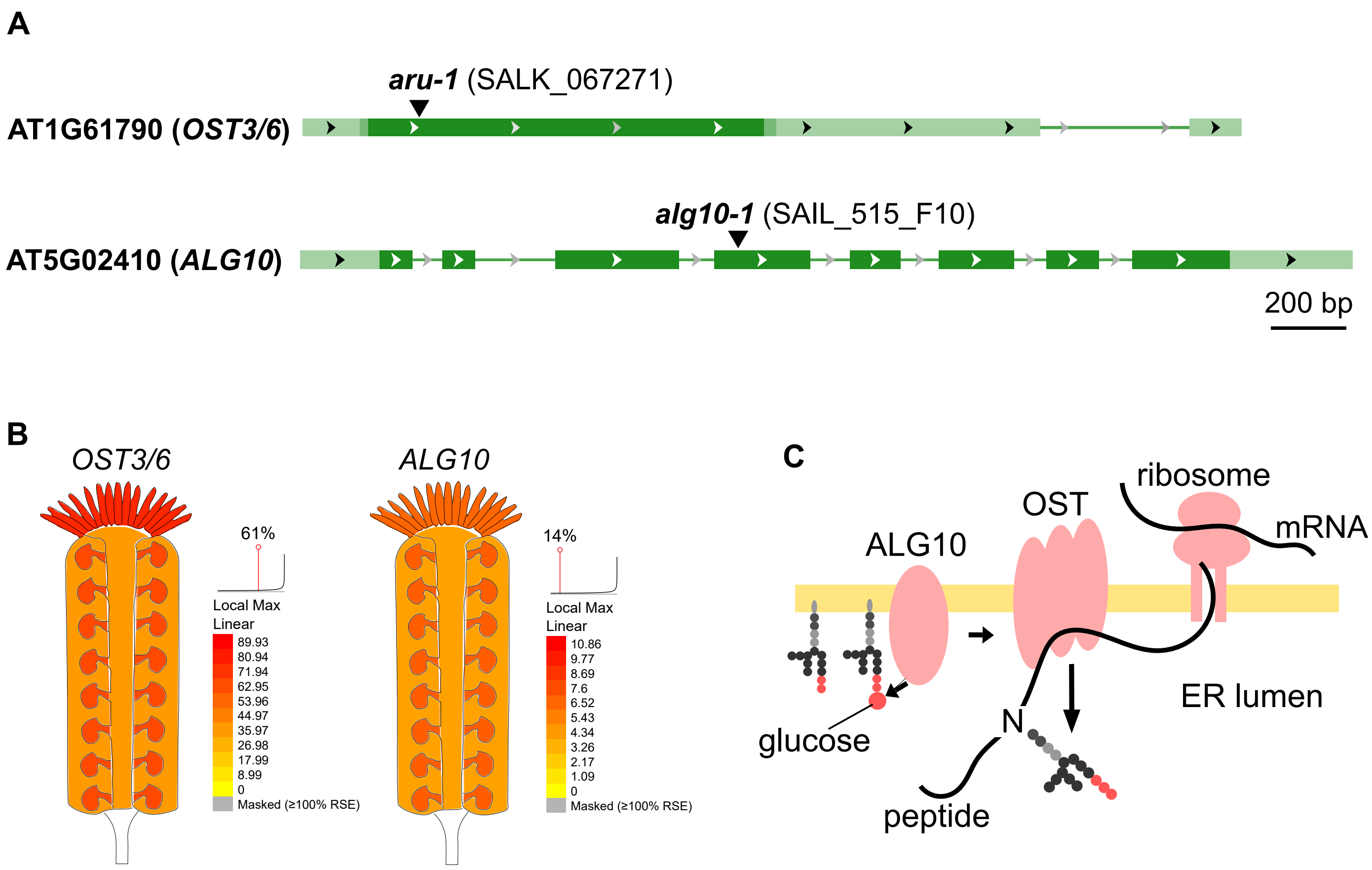


**Supplementary Figure S2** Expression and function of OST3/6 and ALG10. (A) Schematic overview of the structure of the *OST3/6* and *ALG10* genes according to NCBI. The boxes represent mRNA sequences, and the dark green areas represent the coding region. T-DNA insertion sites are also indicated. (B) Expression pattern of *OST3/6* and *ALG10* in pistil tissues according to ePlant (Swanson et al. 2005). The color scale represents the linear expression of genes considering the stigma and ovaries tissues: yellow indicates low expression levels and red indicates high expression levels. (C) Schematic overview of N-glycan biosynthesis by ALG10 and OST.  ALG10 add the third glucose residue to the terminus of the glycan and then OST complex transcend the N-glycan to the nescent peptide chain.

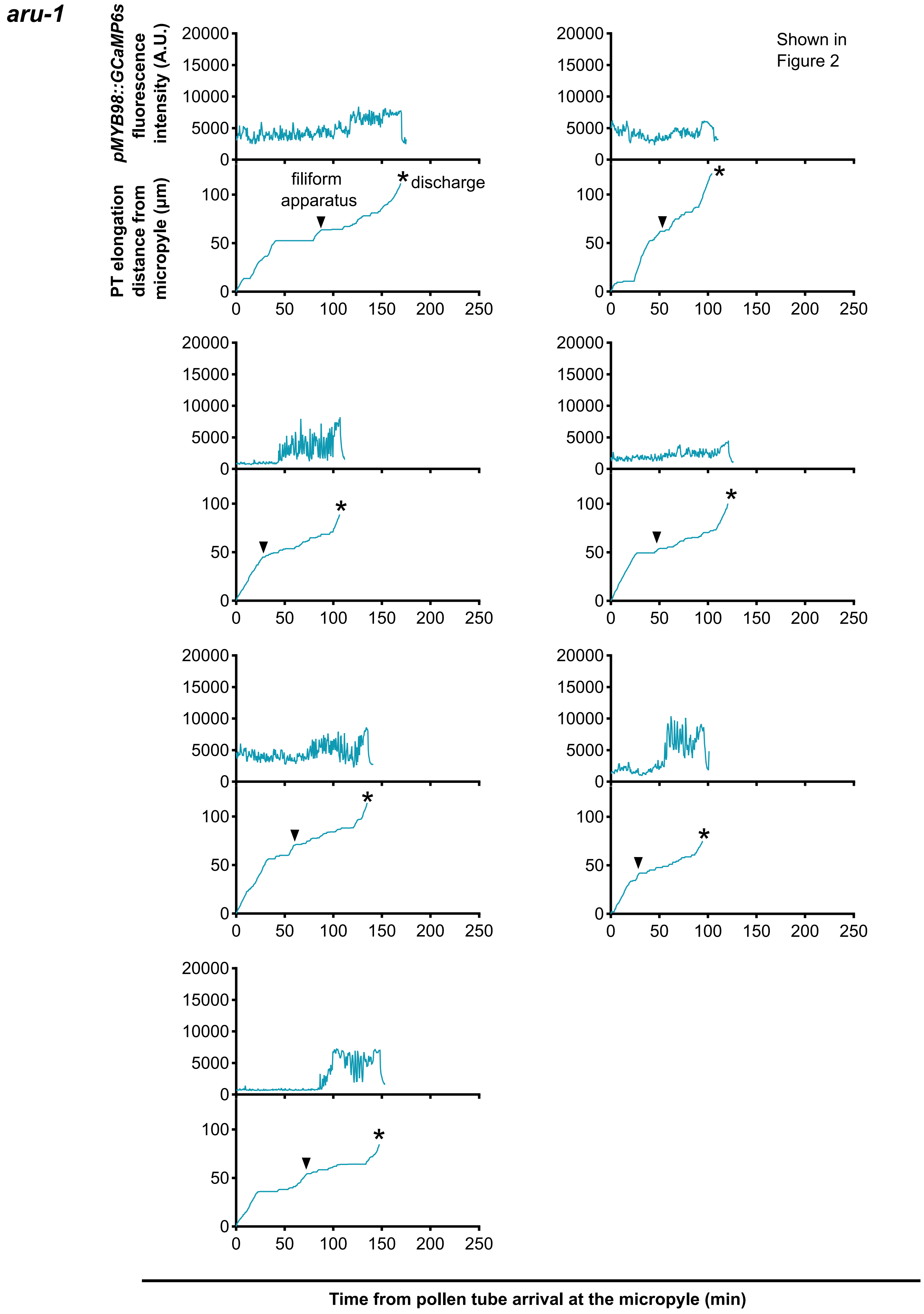


(Continued on next page)

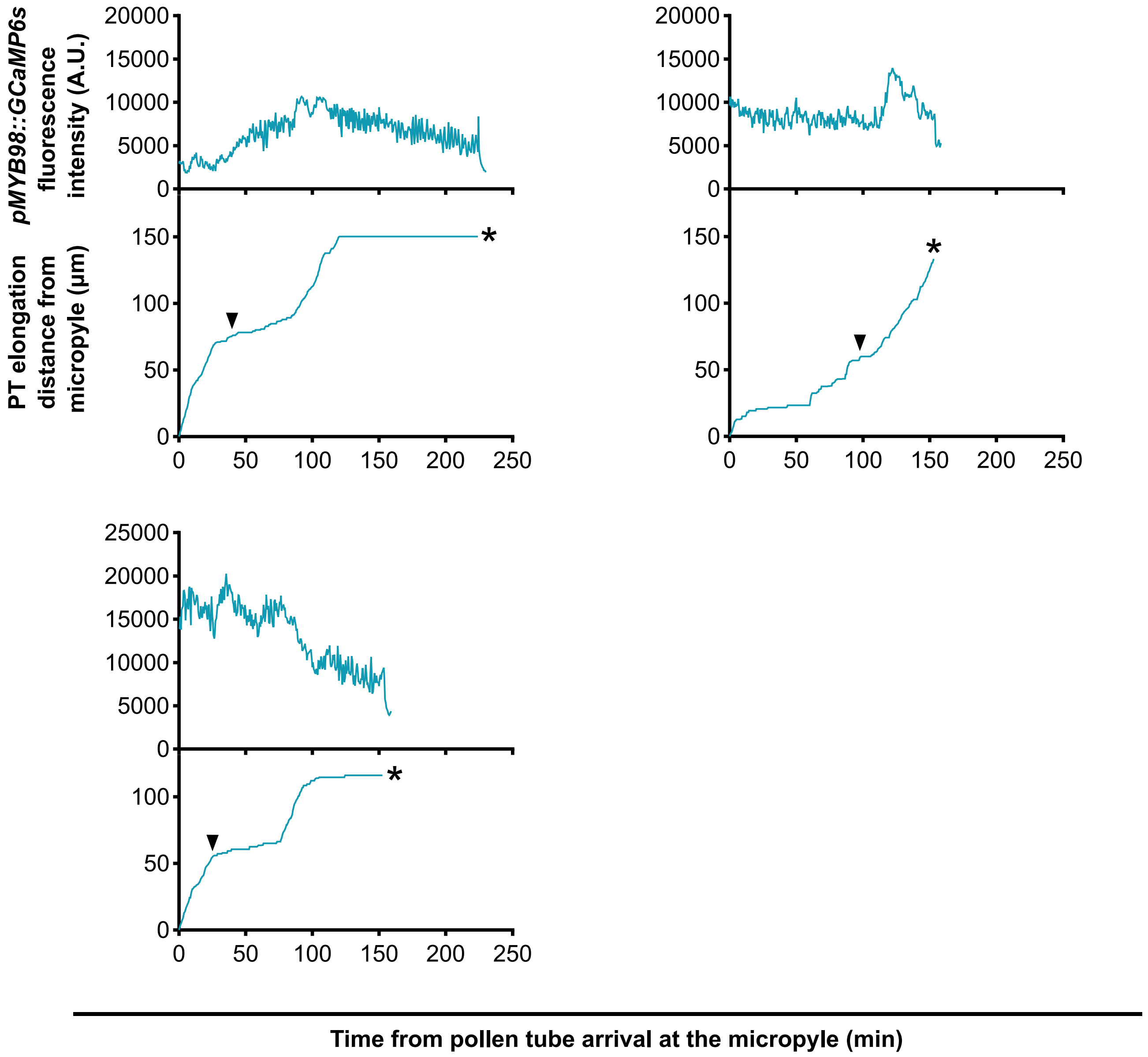


**Supplementary Figure S3** Ten replicate time-lapse calcium ion traces of *pMYB98:GCaMP6s* in the *aru-1* mutant background. The upper graphs show the fluorescence intensity of GCaMP6s in the synergid cells, and the lower graphs show the pollen tube elongation distance from the micropyle. All were pollinated with wild-type pollen. The arrowheads labeled "filiform apparatus" indicate when the pollen tube tip reached the filiform apparatus. Astarisks indicate pollen tube-synergid cell rupture.

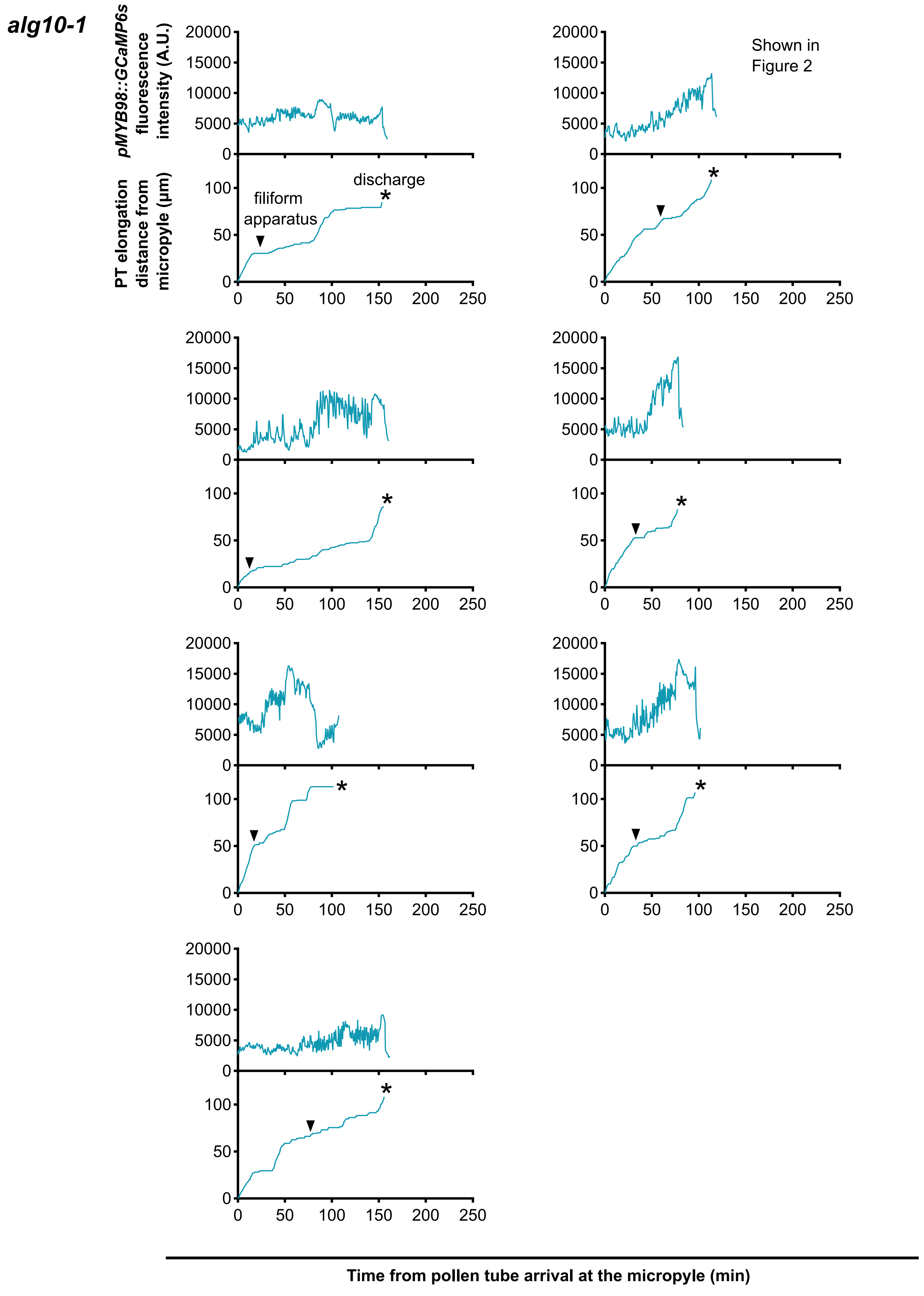


(Continued on next page)

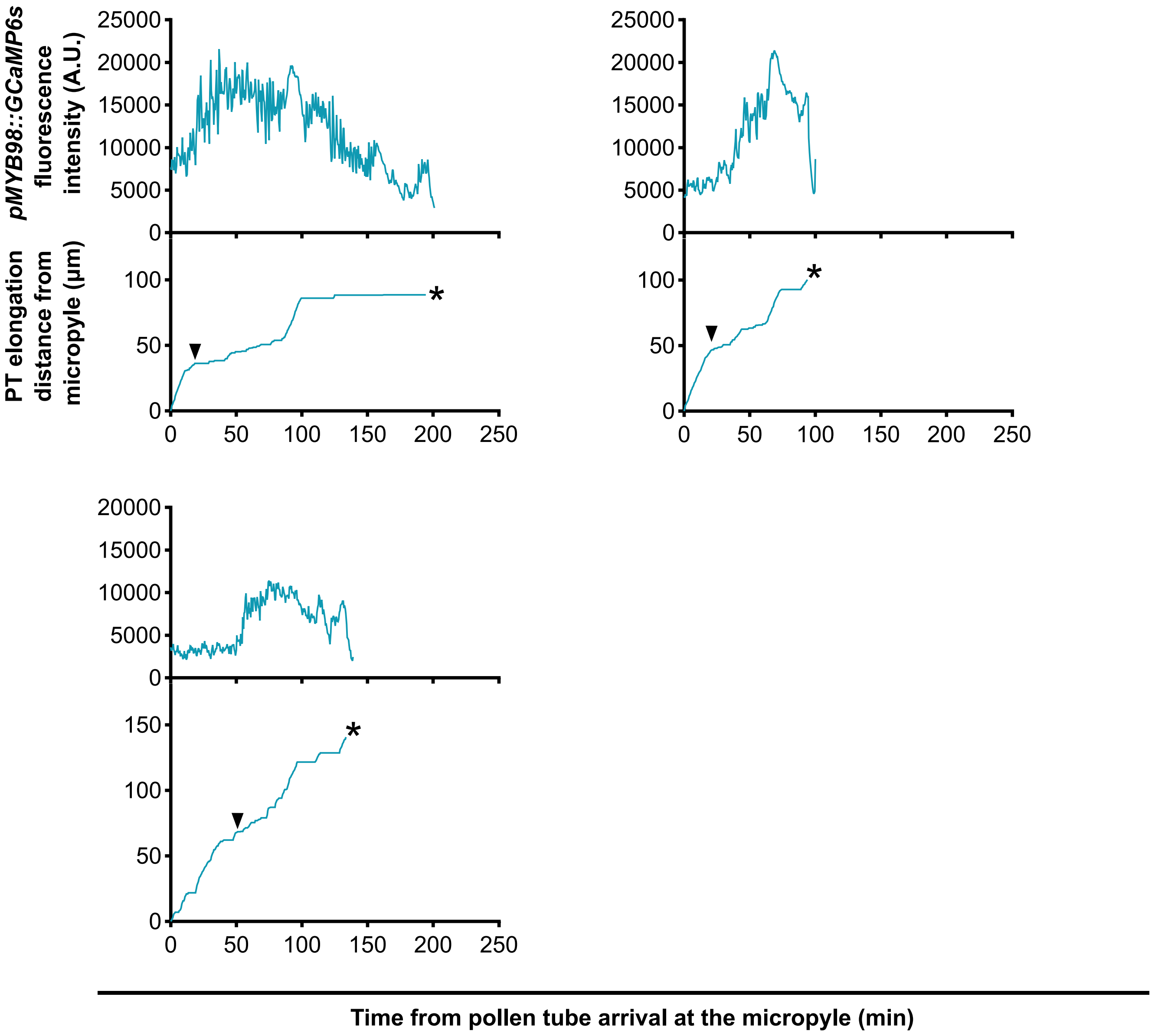


**Supplementary Figure S4** Ten replicate time-lapse calcium ion traces of *pMYB98:GCaMP6s* in the *alg10-1* mutant background. The upper graphs show the fluorescence intensity of GCaMP6s in the synergid cells, and the lower graphs show the pollen tube elongation distance from the micropyle. All were pollinated with wild-type pollen. The arrowheads labeled "filiform apparatus" indicate when the pollen tube tip reached the filiform apparatus. Astarisks indicate pollen tube-synergid cell rupture.

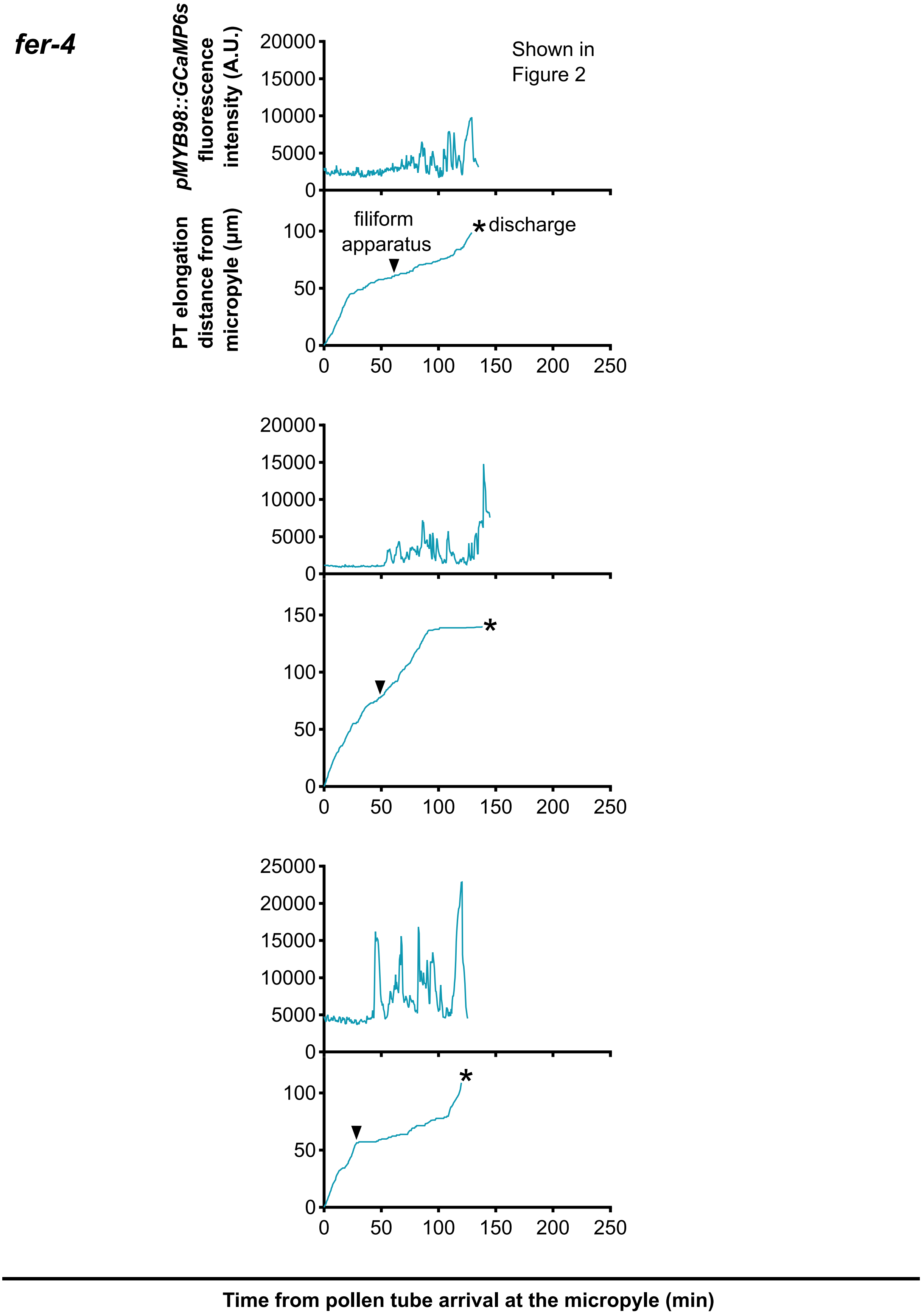


**Supplementary Figure S5** Three replicate time-lapse calcium ion traces of *pMYB98:GCaMP6s* in the *fer-4* mutant background. The upper graphs show the fluorescence intensity of GCaMP6s in the synergid cells, and the lower graphs show the pollen tube elongation distance from the micropyle. All were pollinated with wild-type pollen. The arrowheads labeled "filiform apparatus" indicate when the pollen tube tip reached the filiform apparatus. Astarisks indicate pollen tube-synergid cell rupture.

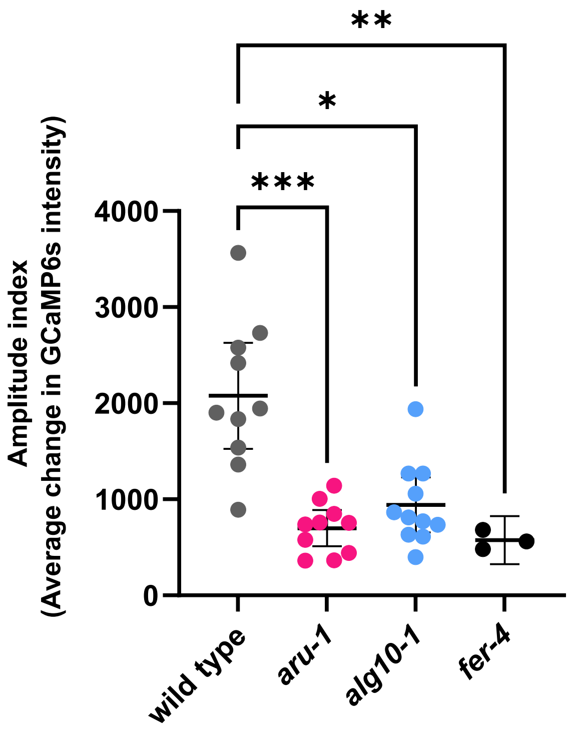


**Supplementary figure S6** Amplitude of synergid calsium oscillations is reduced in *aru-1* and *alg10-1*. Amplitude index of wild type control (n=10), *aru-1* (n=10), *alg10-1* (n=11) and *fer-4* (n=3). We calculated the average change in GCaMP6s intensity per 30 seconds during the period from when the pollen tube reached the filiform apparatus until the receptive synergid cell burst (Phases 1–2). Each point represents an individual ovule and bars represent mean value ± 95% CI. Statistical significance was determined using a Kruskal-Wallis test followed by Dunnett's multiple comparisons test. *p=0.0130; **p=0.0031; ***p=0.0003.

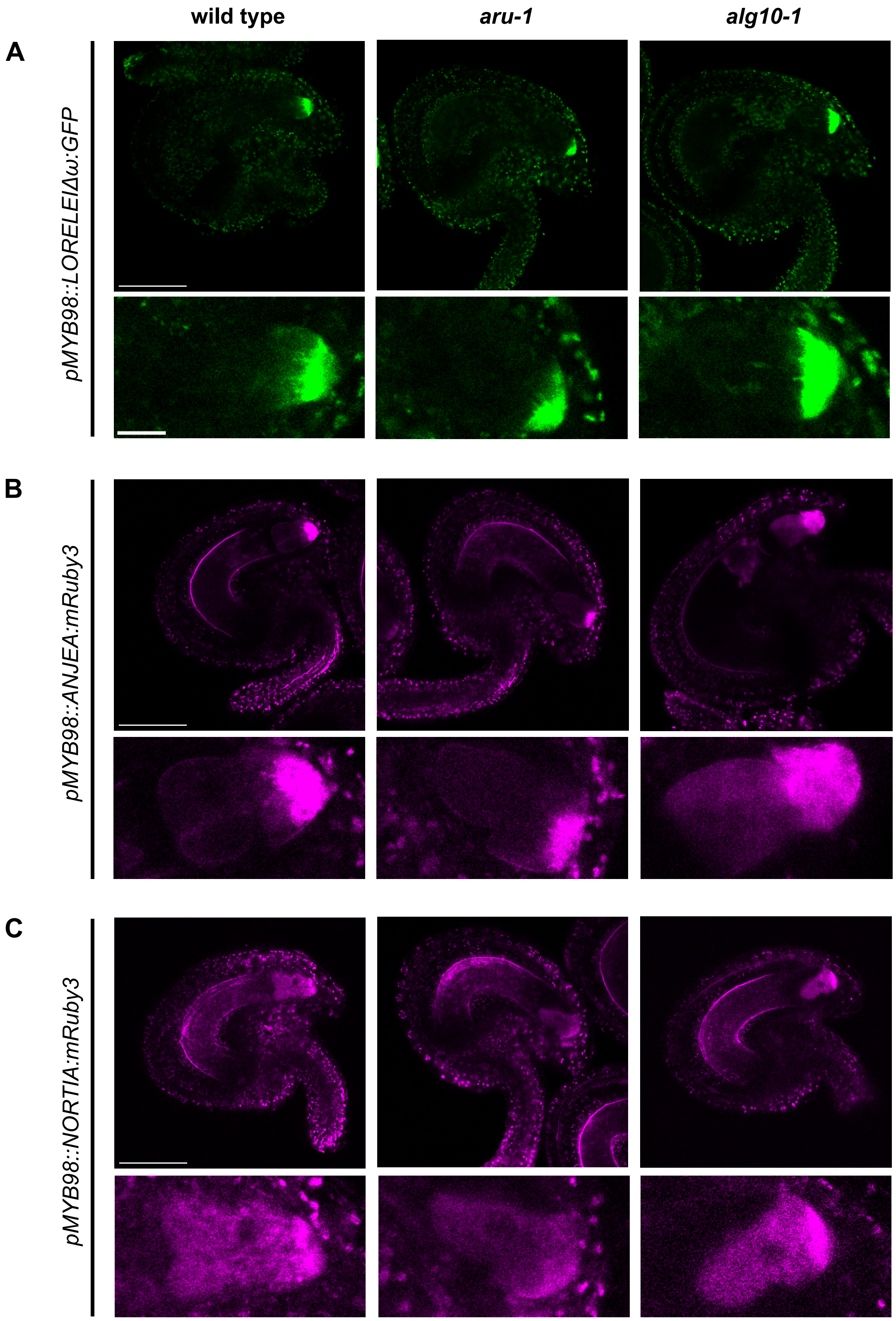


**Supplementary Figure S7** LORELEI, ANJEA and NORTIA exhibit proper localization in the *aru-1* and *alg10-1* mutant synergid cells. (A) LORELEI:GFP localization in the synergid cells of a wild type, *aru-1* and *alg10-1*. (B) ANJEA:mRuby3 localization in the synergid cells of a wild type, *aru-1* and *alg10-1*. (C) NORTIA:mRuby3 localization in the synergid cells of a wild type, *aru-1* and *alg10-1*. (A-C) The upper panels show confocal images of ovules (Scale bar, 50 μm), and the lower panels show close-up images of synergid cells (Scale bar, 10 μm).

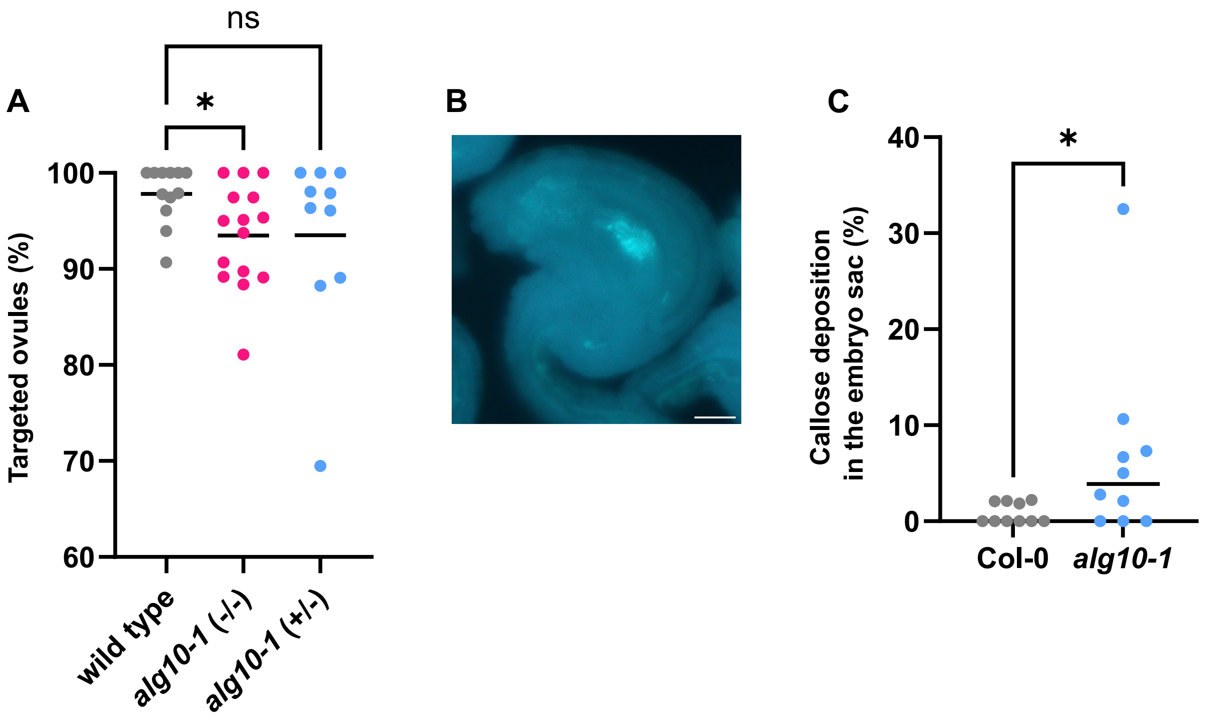


**Supplementary figure S8** *alg10-1* exhibits reduced pollen tube guidance efficiency and a high rate of abnormal ovule development. (A) Percentage of ovules that guided pollen tubes 24 hours after pollination (wild type, n=12 pistils; *alg10-1*(-/-), n=15 pistils; *alg10-1*(+/-), n=10 pistils). Bars represent mean value ± 95% CI. Statistical significance was determined using a Kruskal-Wallis test followed by Dunnett's multiple comparisons test. *p=0.0438. ns not significant. (B) Aniline blue stained *alg10-1* ovule showing abnormal ovule development accompanied by increased callose accumulation. Scale bar, 20 μm. (C) Statistacal analysis of (B). Bar represents mean value. Each point represents an individual pistil (wild type, n=10; *alg10-1*, n=10). Statistical significance was determined using a Mann-Whitney test.
